# Operating Characteristics of the Rank-Based Inverse Normal Transformation for Quantitative Trait Analysis in Genome-Wide Association Studies

**DOI:** 10.1101/635706

**Authors:** Zachary R. McCaw, Jacqueline M. Lane, Richa Saxena, Susan Redline, Xihong Lin

## Abstract

Quantitative traits analyzed in Genome-Wide Association Studies (GWAS) are often non-normally distributed. For such traits, association tests based on standard linear regression are subject to reduced power and inflated type I error in finite samples. Applying the rank-based Inverse Normal Transformation (INT) to non-normally distributed traits has become common practice in GWAS. However, the different variations on INT-based association testing have not been formally defined, and guidance is lacking on when to use which approach. In this paper, we formally define and systematically compare the direct (D-INT) and indirect (I-INT) INT-based association tests. We discuss their assumptions, underlying generative models, and connections. We demonstrate that the relative powers of D-INT and I-INT depend on the underlying data generating process. Since neither approach is uniformly most powerful, we combine them into an adaptive omnibus test (O-INT). O-INT is robust to model misspecification, protects the type I error, and is well powered against a wide range of non-normally distributed traits. Extensive simulations were conducted to examine the finite sample operating characteristics of these tests. Our results demonstrate that, for non-normally distributed traits, INT-based tests outperform the standard untransformed association test (UAT), both in terms of power and type I error rate control. We apply the proposed methods to GWAS of spirometry traits in the UK Biobank. O-INT has been implemented in the `R` package `RNOmni`, which is available on CRAN.

## 1. Introduction

In Genome-Wide Association Studies (GWAS) of continuous (quantitative) traits, the covariate-adjusted genetic effect is typically estimated by linear regression using ordinary least squares (OLS). When the residual distribution is normal, the OLS estimator is normally distributed, consistent, and efficient (Rawlings et al., 1998). However, for many complex traits, including spirometry measurements, the residual distribution is markedly non-normal. An example is peak expiratory flow (PEF), whose residual distribution is skewed and asymmetric even when the outcome is log transformed. When the residual distribution is non-normal, but has mean zero and finite-variance, the OLS estimator remains consistent and asymptotically normal (Cameron and Trivedi, 2005). However, the discrepancy between the asymptotic and finite-sample distributions of the test statistic makes association tests based on the OLS estimator sensitive to the underlying residual distribution (Rawlings et al., 1998). Due to slower convergence of the sampling distribution in the tails, excessive sample sizes *n* ≫ 10^5^ may be required to achieve nominal control of the type I error at the genome-wide significance threshold of *α* = 5*×*10^*−*8^. Moreover, even if the sample is sufficiently sized to protect the type I error, the OLS estimator is no longer efficient when the residual distribution is non-normal (Serfling, 1980). Consequently, OLS-based association tests may lack power for detecting true effects. These limitations of standard association tests in finite samples (failure to control the type I error and poor power) are highlighted in our simulation studies.

The rank-based inverse normal transformation (INT) is commonly applied during GWAS of non-normally distributed traits. INT is a non-parametric mapping that replaces sample quantiles by quantiles from the standard normal distribution. After INT, the marginal distribution of any continuous outcome is asymptotically normal. INT has the effect of symmetrizing and concentrating the residual distribution around zero. Based on the Edgeworth expansion (Lehmann, 1999), convergence of the OLS estimator’s sampling distribution is accelerated when the residual distribution is more nearly normal. Heuristically, INT improves the operating characteristics of standard association testing by increasing residual normality, which in turn allows the sampling distribution of the test statistic to converge more quickly.

We classify INT-based tests into direct and indirect methods. In the direct method (D-INT), INT is applied directly to the phenotype, and the INT-transformed phenotype is regressed on genotype and covariates. Covariates may include age, sex, and adjustments for population structure, such as genetic ancestry principal components (PCs). D-INT has been applied to GWAS of BMI (Scuteri et al., 2007), circulating lipids (Barber et al., 2010), polysomnography signals (Cade et al., 2016), and many quantitative traits in the UK Biobank (Abbott et al., 2017). In the indirect method (I-INT), the phenotype is first regressed on covariates to obtain residuals, then the INT-transformed phenotypic residuals are regressed on genotype, with or without secondary adjustment for population structure. I-INT has been applied to GWAS of gene expression (Emilsson et al., 2008; Consortium et al., 2017), serum metabolites (Kettunen et al., 2012), and spirometry measurements (Repapi et al., 2010). However, the relative performance of D-INT versus I-INT has not been studied in detail. For the non-normal quantitative traits encountered in practice, it is unclear which of the these methods will more robustly control the type I error, and which will provide better power.

As discussed by Beasley and colleagues (Beasley et al., 2009), the question of whether INT-based methods have desirable operating characteristics in the GWAS context has not been critically evaluated. INT of the outcome in a regression model does not guarantee correct model specification. This is because standard linear regression, considered parametrically, requires normality of the residual distribution, not of the marginal distribution of the outcome (Rawlings et al., 1998). Here we systematically study the direct and indirect INT-based association tests, and provide recommendations on how to apply the INT in practice. We begin by formally defining D-INT and I-INT, studying their underlying assumptions and connections. We demonstrate that if the observed trait is generated by a non-linear, rank-preserving transformation of a latent normal trait, then INT provides an approximate inverse of the generative transformation, under the null hypothesis and supposing the covariate effects are small. Moreover, if the mean of the observed trait is linear in covariates but the residual distribution is non-normal, then I-INT is asymptotically exact. Our derivation of I-INT agrees with recent work recommending double adjustment for covariates during INT-based association testing (Sofar et al., 2019).

Through extensive simulations covering the types of residual non-normality often encountered in practice, we compare D-INT and I-INT with the standard untransformed association test (UAT). We find that INT-based tests robustly control the type I error and dominate the UAT in terms of power. However, neither D-INT nor I-INT was uniformly most powerful, and their relative performance depended on the underlying data generating mechanism. Since this is seldom known in practice, we next propose an adaptive omnibus test (O-INT) that synthesizes D-INT and I-INT. O-INT robustly controls the type I error, and across traits is nearly as powerful as the more effective of the component methods. We have implemented all candidate INT-based tests (D-INT, I-INT, and O-INT) in the R package `RNOmni`, which is available on CRAN.

We applied the UAT and the INT-based association tests to GWAS of spirometry traits in the UK Biobank (Sudlow et al., 2015). To demonstrate the power advantage provided by O-INT, we compare the results from the overall analysis (*n* = 292K) with those from a subgroup analysis (*n* = 29K) among asthmatics. All associations identified by O-INT in the asthmatic subgroup were declared significant by UAT in the overall analysis. Hence UAT and O-INT tests agree as to the importance of these loci. However, the more-powerful O-INT test was able to detect them using only a fraction (9.7%) of the sample. In both the asthmatic subgroup and the overall analysis, O-INT realized empirical efficiency and discovery gains over the UAT.

The remainder of this paper is structured as follows. In the methods section, we present all candidate association tests, theoretically study D-INT and I-INT, and propose the INT-based omnibus test (O-INT). In the simulation studies, we present evidence that INT-based association tests robustly control the type I error, whereas the UAT often does not. We demonstrate that INT-based association tests dominate the UAT in terms of power, and show that O-INT is an effective compromise between D-INT and I-INT. In the data application, we compare the performance of all candidate association tests for GWAS of spirometry traits from the UK Biobank. We conclude with a discussion of the implications of our findings for quantitative trait GWAS.

## 2. Statistical Methods

### 2.1 Setting

For each of *n* independent subjects, the following data are observed: a continuous (quantitative) phenotype *Y*_*i*_, genotype *g*_*i*_ at the locus of interest, and a *p*×1 vector ***x***_*i*_ = (*x*_*i*,1_, … , *x*_*i,p*_) of covariates. In our data application, the phenotype *Y*_*i*_ is a spirometry measurement, while the covariates include an intercept, age, sex, and genetic principal components (PCs). Let ***y*** = (*Y*_1_, … , *Y*_*n*_) denote the *n* × 1 sample phenotype vector, ***g*** the *n* × 1 genotype vector, and ***X*** the *n* × *p* covariate design matrix.

### 2.2 Untransformed Association Test

The untransformed association test (UAT) is derived from the normal linear model:

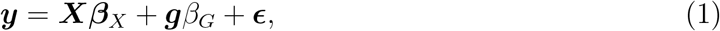

where ***ϵ*** ∼ *N*(**0**, *σ*^2^***I***) is an *n* × 1 residual vector, *β*_*G*_ is the genetic effect, and ***β***_*X*_ is the covariate effect. Define the error projection ***P***_*X*_ = ***I*** − ***X***(***X′X***)^*−*1^***X′***, and the *phenotypic residual **e***_*Y*_ = ***P***_*X*_***y***, which is the residual after regressing ***y*** on ***X***, or the projection of ***y*** onto the orthogonal complement of the column space of ***X***. The efficient score for *β*_*G*_ is 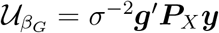, and the score statistic assessing *H*_0_: *β*_*G*_ = 0 is:

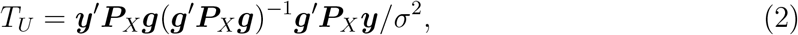

which follows a 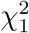 distribution. Under *H*_0_, an unbiased estimate of the residual variance is given by 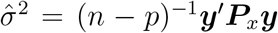. The Wald statistic for assessing *H*_0_: *β*_*G*_ = 0 takes the same form as (2), save the residual variance is estimated as 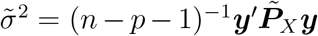, where 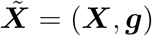 and 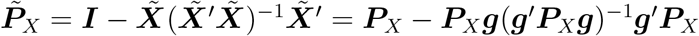.

If the normal residual assumption is relaxed to allow for an arbitrary distribution with mean zero and finite variance, then *T*_*U*_ still follows an asymptotic 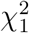 distribution. Although (2) is eventually valid for any continuous trait with constant residual variance, when the residual distribution exhibits excess skew or kurtosis, the sample size required for valid inference at *α* = 5 *×* 10^*−*8^ may become impractically large. Moreover, as we will show, even in samples of sufficient size for valid inference, the UAT is generally less powerful than INT-based tests.

### 2.3 Rank-Based Inverse Normal Transformation

INT is a non-parametric mapping applicable to observations from any absolutely continuous distribution. The process may be decomposed into two steps. In the first, the observations are replaced by their fractional ranks. This is equivalent to transforming the observations by their empirical cumulative distribution function (ECDF) 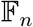. If *W* is any continuous random variable with CDF *F*_*W*_, then the transformed random variable *U* = *F*_*W*_ (*W*) is uniformly distributed (Casella and Berger, 2002). Since the empirical process 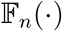 converges uniformly to the CDF *F*_*W*_, in independent and identically distributed samples of sufficient size 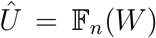 is uniformly distributed (van der Vaart, 1998). After transformation by 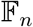, the observations reside on the probability scale. In the next step, these probabilities are mapped to Z-scores using the *probit* function Φ^*−*1^. If *U* is uniformly distributed, then *Z* = Φ^*−*1^(*U*) follows the standard normal distribution. Consequently, in large samples, 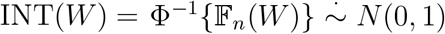, regardless of the initial distribution *F*_*W*_.

In practice, an offset is introduced to ensure that all fractional ranks are strictly between zero and one, which in turn guarantees that all Z-scores are finite. Suppose that *W*_*i*_ is observed for each of *n* independent subjects. The modified INT is:

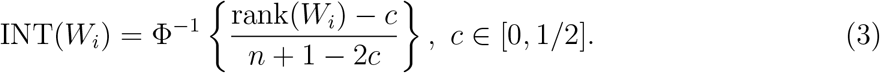

Hereafter we adopt the conventional *Blom* offset of *c* = 3*/*8 (Beasley et al., 2009). Since other choices for *c* lead to Z-scores that are nearly linear transformations of one another, the choice of offset is considered immaterial.

### 2.4 Direct Inverse Normal Transformation (D-INT)

In direct INT (D-INT), the INT-transformed phenotype is regressed on genotype and covariates according to the association model:

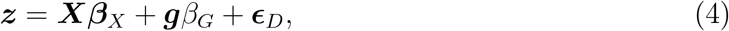

where ***z*** = INT(***y***) is the INT-transformed phenotype, and 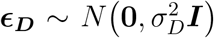. Model (4) is immediately comparable with model (1), the only difference being replacement of ***y*** by ***z***. Thus, the D-INT score statistic for assessing *H*_0_: *β*_*G*_ = 0 is:

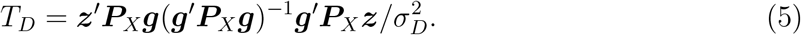

A *p*-value is assigned with reference to the 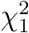 distribution. The score statistic estimates the residual variance as 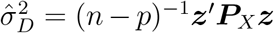, whereas the Wald statistic estimates the residual variance as 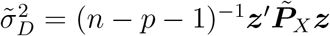.

D-INT is adapted for data generating processes (DGPs) of the form:

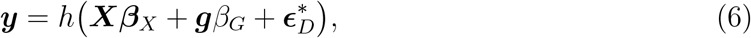

where 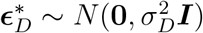, and *h*(*⋅*) is a rank-preserving transformation. An example is the lognormal phenotype, for which *h*(*t*) = exp(*t*). When *h*(*⋅*) is non-linear, the regression function *E*(*Y*_*i*_|***x***_*i*_, *g*_*i*_) is non-linear in its parameters, and the residuals 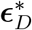 have non-additive effects. However, there exists a transformed scale on which the mean model is linear and has additive normal residuals, namely: *h*^−1^(***y***) = ***Xβ***_*X*_ + ***g**β*_*G*_ + ***ϵ***_*D*_. Thus, under *H*_0_: *β*_*G*_ = 0, the efficient score for *β*_*G*_ is 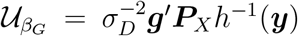. Since *h* ^−1^(*⋅*) is seldom known, D-INT makes the approximation 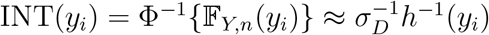 where 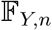 is the *marginal* ECDF of the phenotype *Y*_*i*_.

To justify this approximation, observe that under model (6), the *conditional* distribution of the transformed-scale residual 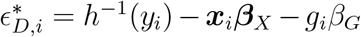 is 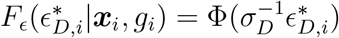.

The marginal and conditional CDFs of *Y*_*i*_ are related via:

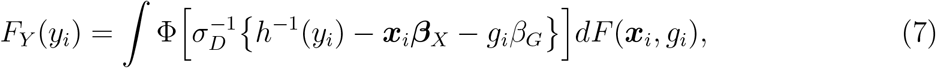

where *F* (***x***_*i*_, *g*_*i*_) is the joint CDF of ***x***_*i*_ and *g*_*i*_. The empirical counterpart to (7) is:

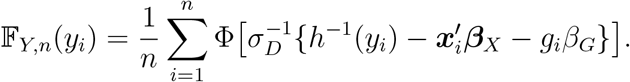

Under the complete null *H*_0_: (***β***_*X*_ = 0) *and* (*β*_*G*_ = 0), 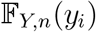 converges to 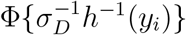, such that the D-INT approximation 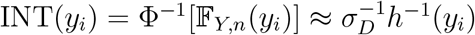 is asymptotically exact. Under the standard *H*_0_: *β*_*G*_ = 0, the approximation is accurate when ***β***_*X*_ ≈ **0**.

### 2.5 Indirect Inverse Normal Transformation (I-INT)

In indirect INT (I-INT), the phenotype is first regressed on covariates to obtain residuals, then the INT-transformed phenotypic residuals are regressed on genotype. Specifically, I-INT is based on the association model:

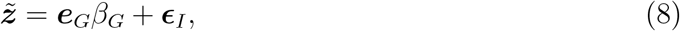

where 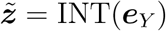 is the INT-transformed phenotypic residual (i.e. ***e***_*Y*_ = ***P***_*X*_***y***), ***e***_*G*_ = ***P***_*X*_***g*** is the *genotypic residual*, which is the residual after regressing ***g*** on ***X***, and 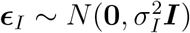. The I-INT score statistic for assessing *H*_0_: *β*_*G*_ = 0 takes the form:

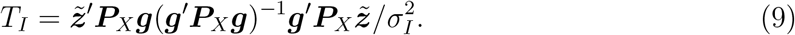

A *p*-value is assigned with reference to the 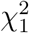 distribution. The score statistic estimates the residual variance as 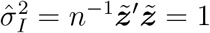, while the Wald statistic estimates the residual variance as 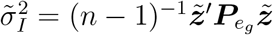, where 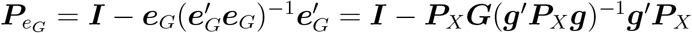.

Since INT is a non-linear transformation, the INT transformed phenotypic residuals 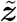 are no longer orthogonal to the covariates. That is, the correlation between 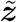 and the columns of ***X*** is non-zero. Consequently, secondary adjustment for covariates has been recommended Sofar et al. (2019), as in the association model:

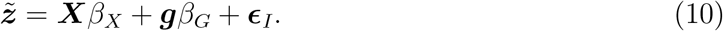

We demonstrate that the score statistic from model (10), which adjusts twice for covariates, is equivalent to the score statistic from model (8), which instead adjusts for genotypic residuals.

I-INT is adapted for a DGP of the form:

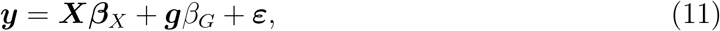

where the residuals ***ε*** ∼ *f*_*ε*_(*⋅*) follow an arbitrary continuous distribution with mean zero and constant finite variance. Under (11), 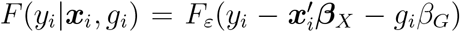, such that under the complete null *H*_0_: (***β***_*X*_ = 0) *and* (*β*_*G*_ = 0), the DGPs in (11) and (6) are equivalent.

To motivate I-INT, we begin by showing that the efficient score for *β*_*G*_ from model (11) is consistently estimated by the score for *β*_*G*_ from the model ***e***_*Y*_ = ***e***_*G*_*β*_*G*_ + ***ε***. Observe that, for any *f*_*ε*_, the ordinary least squares estimator 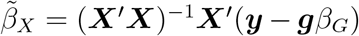 remains consistent for ***β***_*X*_. Thus, the profile log likelihood from (11) is consistently estimated by:

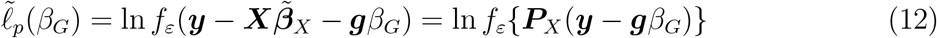

Letting 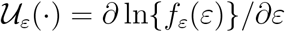, the efficient score for *β*_*G*_ from (11) is consistently estimated by the gradient of (12), which is 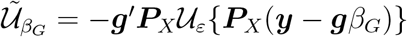.

Now consider the following model for the phenotypic residual:

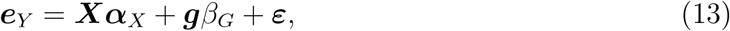

where ***ε*** is distributed as before. The profile log likelihood for *β*_*G*_ in (13), with ***α***_*X*_ evaluated at the consistent estimator 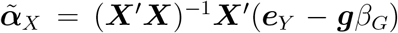, coincides with the profile log likelihood in (12). Moreover, the log likelihood for *β*_*G*_ from the following model, which relates ***e***_*Y*_ to the genotypic residual ***e***_*G*_, is also identical to (12):

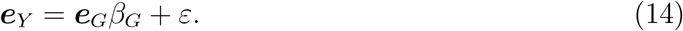

Thus, the score for *β*_*G*_ from (14) is consistent for the efficient score for *β*_*G*_ from (11).

Now, under *H*_0_: *β*_*G*_ = 0, model (14) and model (13) with ***α***_*X*_ evaluated at its least squares estimate 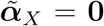, both reduce to ***e***_*Y*_ = ***ε***. Model (14), together with the observation that 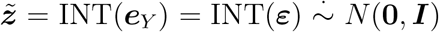, motivate the I-INT association model in (8). Moreover, under *H*_0_: *β*_*G*_ = 0, the distributional assumption in (8) is asymptotically exact, with 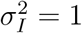.

### 2.6 Omnibus Inverse Normal Transformation Test (O-INT)

As shown in the simulation studies, both D-INT and I-INT robustly controlled the type I error. However, neither D-INT nor I-INT was uniformly most powerful. We therefore propose combining the two approaches into a robust and powerful omnibus test. The omnibus statistic is constructed using the method of Cauchy aggregation, in which the p-values from dependent hypothesis tests are converted to standard Cauchy random deviates and then combined (Liu and Xie, 2019; Liu et al., 2019). Cauchy aggregation is preferred to classic approaches for combining p-values, such as Fisher’s method (Fisher, 1934), since analytical expression are available for the finite-sample distribution of a Cauchy combination of dependent p-values. Let *p*_*D*_ and *p*_*I*_ denote the p-values from D-INT and I-INT. Define the O-INT statistic as:

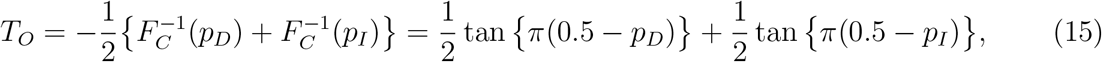

where 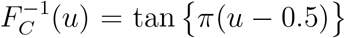 is the inverse CDF of the standard Cauchy distribution. Under *H*_0_: *β*_*G*_ = 0, *p*_*D*_ and *p*_*I*_ are each uniformly distributed, such that 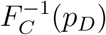 and 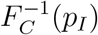 are standard Cauchy. Since the Cauchy distribution is symmetric and closed with respect to convolution, the omnibus statistic 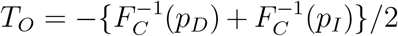 is again standard Cauchy in the tail (Liu and Xie, 2019; Liu et al., 2019), even though *p*_*D*_ and *p*_*I*_ are in general positively correlated. The p-value of the O-INT statistic (15) with observed value *t*_*O*_ is:

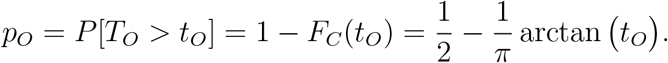

## 3. Simulation Studies

### 3.1 Simulation Methods

Extensive simulations were conducted to evaluate the type I error and power of the UAT and INT-based association tests (D-INT, I-INT, O-INT). Genotypes exhibiting linkage disequilibrium were randomly sampled from unrelated subjects in the UK Biobank. The genotypes were additively coded, assuming values *g*_*i*_ ∈ {0, 1, 2}. Simulated covariates included age and sex. Age was drawn from a gamma distribution with mean 50 and variance 10, and sex was drawn independently from a Bernoulli distribution with proportion 1/2. To emulate population structure, the top 3 PCs of the empirical genetic relatedness matrix were included as covariates. These correspond to the leading 3 left singular vectors from the subject by variant genotype matrix.

For type I error simulations, a subject-specific linear predictor *η*_*i*_ was generated as 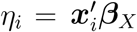, where ***x***_*i*_ included an intercept, age, sex, and 3 genetic ancestry PCs. Regression coefficients were selected such that the proportion of total phenotypic variation explained (PVE) by age and sex was 20%, and the PVE by PCs was 5%. For power simulations, the linear predictor included a contribution from genotype. The PVE by genotype or heritability, defined as *h*^2^ = Var(*g*_*i*_*β*_*G*_)/Var(*Y*_*i*_), ranged between 0.1% and 1.0%.

Phenotypes were generated either from models with additive residuals, as in *y*_*i*_ = *η*_*i*_ + *ϵ*_*i*_, or from non-linear transformations of such models, as in *y*_*i*_ = *h*(*η*_*i*_ + *ϵ*_*i*_). Here, we report on four representative traits: three with additive residuals, and one with multiplicative residuals. The additive models were: (1) a reference trait, with *N* (0, 1) residuals; (2) a skewed trait, with 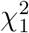 residuals; and (3) a kurtotic trait, with *t*_3_ residuals. In all cases, the residual distribution was centered and scaled to have mean zero and unit variance. For the multiplicative model, a log-normal phenotype was generated by exponentiating a latent normal trait: *y*_*i*_ = exp(*η*_*i*_ + *ϵ*_*i*_), where *ϵ*_*i*_ ∼ *N* (0, 1).

### 3.2 Type I Error Simulations

A total of *R* = 10^8^ simulation replicates were performed under *H*_0_: *β*_*G*_ = 0 at samples size of *n* ∈ {10^3^, 10^4^, 10^5^}. On each simulation replicate, the four phenotypes (normal, skewed, kurtotic, log-normal) were generated independently and tested for association with genotype by each of the four association methods (UAT, D-INT, I-INT, O-INT).

The uniform QQ plots in Figure 1 summarize the distribution of association p-values at sample size *n* = 10^3^ for each combination of phenotype (row) and association test (column). All association tests performed well against the normal phenotype (row 1), providing uniformly distributed p-values. UAT (column 1) exhibited inflated type I error against all non-normal phenotypes, although inflation attenuated with increasing sample size (Web Figures S1-2). Inflation was most severe for the log-normal phenotype (row 4), likely because the standard linear model is misspecified when the residuals have multiplicative rather than additive effects. However, inflation was still present for the skewed 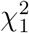 (row 2) and kurtotic *t*_3_ (row 3) phenotypes, for which UAT is correctly specified. In contrast, by sample size *n* = 10^3^ the INT-based tests provided uniformly distributed p-values when applied to non-normal phenotypes. Although the modeling assumptions underlying D-INT were not met for the skewed or kurtotic phenotypes, D-INT maintained the type I error across all scenarios. I-INT exhibited slight deflation against the log-normal phenotype, for which its modeling assumptions were not met. This deflation ameliorated with increasing sample size. O-INT performed well under all scenarios.

**Figure 1.**
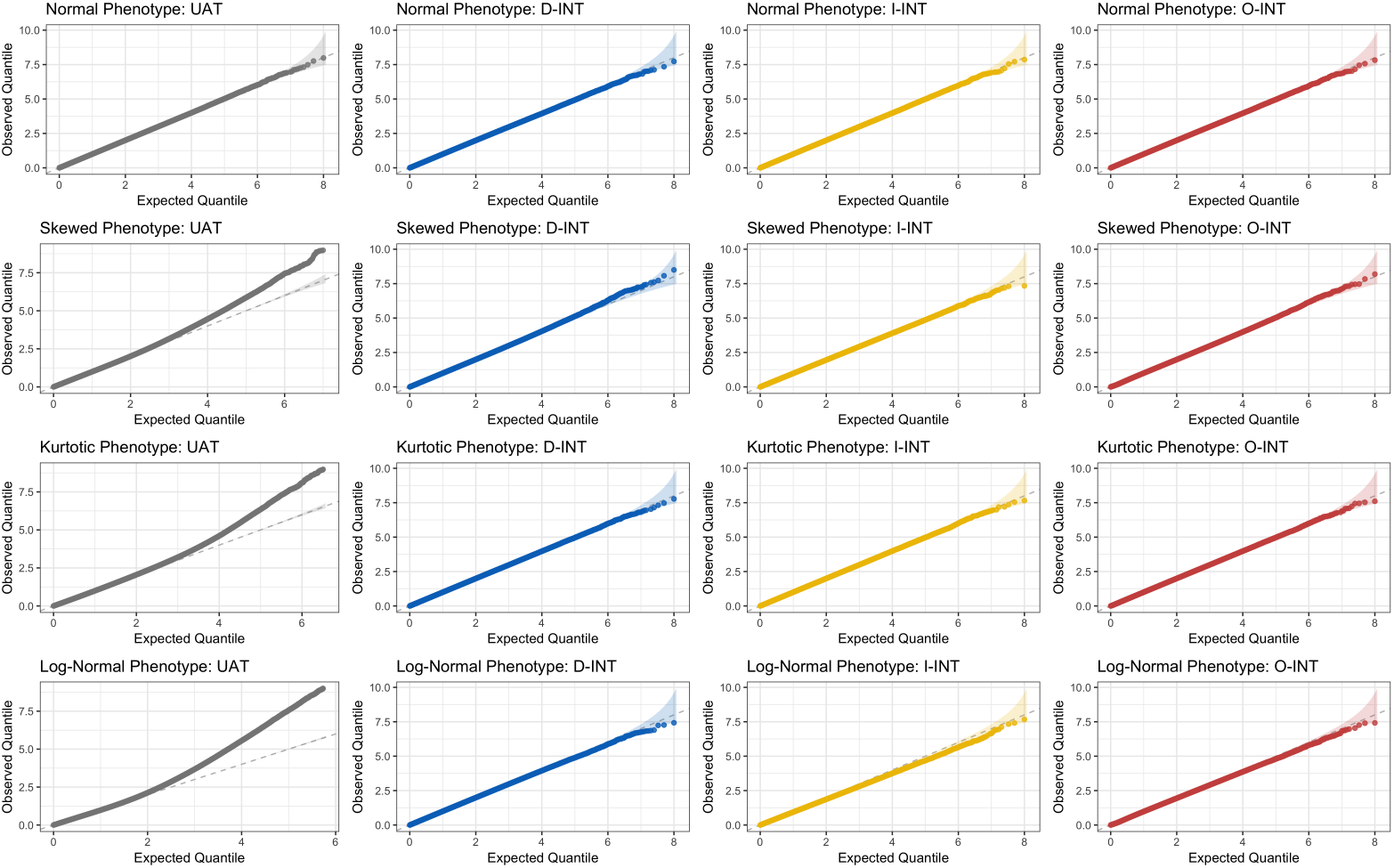
Distribution of Association p-values Under the Null at Sample Size *n* = 10^3^ across *R* = 10^8^ Simulation Replicates. Rows correspond to different phenotype distributions. The first phenotype has normal residuals; the second has 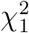 residuals; the third phenotype has *t*_3_ residuals; and the log of the fourth phenotype has normal residuals. Columns correspond to different association tests. The first is the untransformed association test (UAT), the second is the direct INT (D-INT), the third is indirect INT (I-INT), and the fourth column is omnibus INT (O-INT). Note that this figure appears in color in the electronic version of this article.

Type I error estimates at *α* = 10^*−*6^ and sample sizes *n* ∈ {10^3^, 10^4^, 10^5^} are presented in Table 1. For all non-normal phenotypes, UAT had substantially inflated type I error at sample size *n* = 10^3^. This includes the skewed 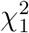 and kurtotic *t*_3_ phenotypes, for which UAT should provide asymptotically valid inference. Although the type I error approached its nominal level with increasing sample size, for the kurtotic and log-normal phenotypes the UAT still exhibited excess type I error at *n* = 10^5^. For the non-normal phenotypes, D-INT generally provided nearly the nominal type I error, while I-INT was slightly conservative. However, this does not imply I-INT is less powerful for these phenotypes (see power simulations). For all phenotypes and sample sizes considered, the omnibus test provided nominal control of the type I error.

**Table 1.** Empirical Type I Error (×10^6^) at α = 10^*−*6^ across R = 10^8^ Simulation Replicates. Size simulations were conducted under the H_0_: *β*_*G*_ = 0 at sample sizes ranging from n = 10^3^ to n = 10^5^. The following association tests were evaluated: the untransformed association test (UAT), direct INT (D-INT), indirect INT (I-INT), and omnibus INT(O-INT). Each test was applied to a normal phenotype, a skewed phenotype with 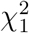 residuals, a kurtotic phenotype with t_3_ residuals, and a phenotype whose log had normal residuals.

| Phenotype | Test | Sample Size |  |  |
| --- | --- | --- | --- | --- |
| | | $n = 10^3$ | $n = 10^4$ | $n = 10^5$ |
| Normal | UAT | 1.04 | 0.93 | 1.03 |
| Normal | D-INT | 0.84 | 0.87 | 1.02 |
| Normal | I-INT | 0.97 | 0.93 | 1.00 |
| Normal | O-INT | 0.91 | 0.93 | 0.99 |
| Skewed | UAT | 8.03 | 1.87 | 1.43 |
| Skewed | D-INT | 1.20 | 1.10 | 1.05 |
| Skewed | I-INT | 0.67 | 0.84 | 0.89 |
| Skewed | O-INT | 1.10 | 1.01 | 0.98 |
| Kurtotic | UAT | 15.89 | 5.54 | 3.12 |
| Kurtotic | D-INT | 0.94 | 0.91 | 0.95 |
| Kurtotic | I-INT | 1.00 | 0.88 | 1.00 |
| Kurtotic | O-INT | 0.96 | 0.90 | 0.97 |
| Log-Normal | UAT | 59.34 | 11.01 | 7.52 |
| Log-Normal | D-INT | 0.74 | 1.02 | 1.02 |
| Log-Normal | I-INT | 0.40 | 0.40 | 0.43 |
| Log-Normal | O-INT | 0.55 | 0.74 | 0.76 |

### 3.3 Power Simulations

At each heritability *h*^2^ ∈ {0.1, 0.2, … , 1.0}%, a total of *R* = 10^6^ power simulations were performed at sample size *n* = 10^3^. On each replicate, a single randomly selected locus served as the causal locus. As before, the phenotypes were generated independently and tested for association with genotype by each of the candidate association methods. Power is considered even for the UAT, which did not consistently control the type I error, because this approach is still often applied in practice.

Power curves at *α* = 10^*−*6^ are presented in Figure 2. Relative efficiency (RE) curves, comarping the INT-based tests with UAT, are presented in Web Figure S3. RE was calculated as the ratio of the 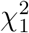 non-centrality parameters. This metric has the advantage of not depending on either *α* level or sample size *n*. For the normal phenotype, the UAT is theoretically most powerful. However, the INT-based tests were fully efficient, achieving a relative efficiency of one. Despite having inflated type I error under the null hypothesis, UAT was consistently least powerful for detecting true associations with the non-normal phenotypes. Since the relative efficiencies of the INT-based tests always exceeded one for the non-normal phenotypes, this conclusion is expected to extend across significance levels and sample sizes. For the log normal phenotype, D-INT was most powerful, achieving twice the efficiency of the UAT. For this phenotype, the log transform is theoretically optimal. Since the log transform maps the log normal phenotype to a normal phenotype, the power of the log transform against the log normal phenotype is identical to the power of the UAT against the normal phenotype. Comparing the power of D-INT against the log normal phenotype with that of UAT against the normal phenotype, we observe that D-INT attains optimal power. For the skewed 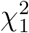 phenotype, I-INT was most powerful, achieving over 5 times the efficiency of the UAT, while D-INT was twice as efficient. For the kurtotic *t*_3_ phenotype, the efficiency gains provided by the INT-based tests were more modest yet still noteworthy, at around 55% for the I-INT and 35% for D-INT.

**Figure 2.**
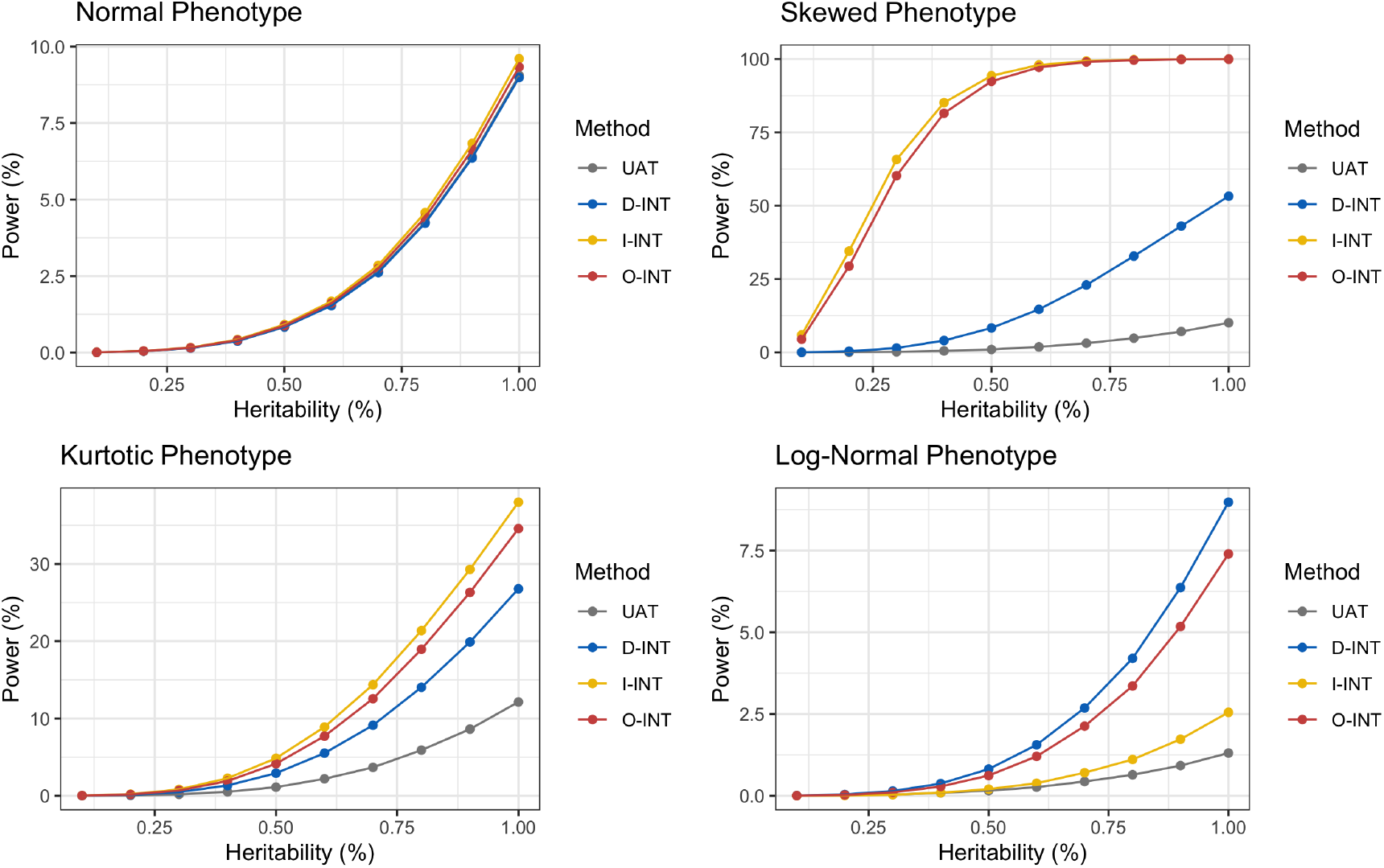
Power Curves at *α* = 10^*−*6^ and Sample Size *n* = 10^3^ across *R* = 10^6^ Simulation Replicates. Simulations were conducted at heritabilities ranging from 0.1% and 1.0%. Gray is the untransformed association test (UAT), blue is direct INT (D-INT), yellow the indirect INT (I-INT), red is omnibus INT (O-INT). Each panel corresponds to a different phenotype. The first phenotype has normal residuals; the second has 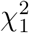 residuals; the third phenotype has *t*_3_ residuals; and the log of the fourth phenotype has normal residuals. Note that this figure appears in color in the electronic version of this article.

By synthesizing D-INT and I-INT, O-INT aims to provide a test that is well powered across the residual distributions encountered in practice. As a compromise between complementary methods, the power and RE of O-INT were intermediate to those of D-INT and I-INT. However, for all phenotypes studies, O-INT performed comparably to the more efficient of D-INT and I-INT. Thus, O-INT achieves robustness to the underlying data generating mechanism with little to no loss of efficiency. In addition, we compared INT-based testing with the non-parametric Kruskal-Wallis (KW) test (Kruskal and Wallis, 1952). Unlike regression-based association tests, adjusting for covariates in the KW test is not straightforward. Yet even in the absence of covariates, INT-based testing was more powerful than the KW test.

## 4. Application to UK Biobank

### 4.1 Application Methods

We conducted GWAS of spirometry phenotypes within the UK Biobank (UKB). To mitigate confounding due to population structure, our study population was restricted to unrelated subjects of white, British ancestry. The phenotypes were forced expiratory volume in 1 second (FEV1), forced vital capacity (FVC), the FEV1 to FVC ratio (FEV1/FVC), and the logarithm of peak expiratory flow (lnPEF). Our analyses focused on 360,761 additively coded and directly genotyped, as opposed to imputed, autosomal SNPs, with sample minor allele frequencies (MAFs) exceeding 5%, and a per locus missingness rates of less than 10%. Covariates included an intercept, age, sex, BMI, two orthogonal polynomials in height, genotyping array, and 20 genetic PCs. Each locus was tested individually for association with the four spirometry phenotypes (FEV1, FVC, FEV1/FVC, lnPEF). The results were greedily ‘clumped’ in PLINK (Purcell et al., 2007) using a 1000 kb radius and an *r*^2^ threshold of 0.2. The overall analysis consisted of *n* = 292K subjects that met all inclusion criteria. A subgroup analyses was conducted among subjects with physician diagnosed asthma (*n* = 29K).

### 4.2 Empirical Type I Error

LD score regression (LDSC) was performed to assess inflation of the association test statistics due to confounding bias (Bulik-Sullivan et al., 2015). Briefly, in LDSC the test statistic for each locus is regressed on a local measure of linkage disequilibrium. An intercept exceeding one suggests inflation, whereas an intercept falling below one suggests deflation. The results from applying LDSC in the overall sample and in the asthmatic subgroup are presented in Web Tables S4-5. Overall, there was no evidence of residual confounding due to population structure. Therefore, under the null hypothesis of no genetic effects, the marginal distribution of each spirometry trait is expected to be independent of genotype.

The empirical type I error of the association tests was assessed via a permutation analysis. Genotypes and phenotypes were first regressed on covariates to obtain residuals, then the genotypic residuals were permuted. Regressing out the effects of covariates accounts for potential confounding of the genotype-phenotype relationship. Permuting the genotypic residuals breaks the association between genotype and the spirometry traits, thereby imposing the null hypothesis of no genetic effect. Uniform QQ plots for the association p-values after permutation are presented in Web Figure S7. For all association methods (columns) and spirometry traits (rows), the p-values were uniformly distributions, suggesting nominal control of the type I error for the observed residual distributions.

### 4.3 Empirical Discovery and Efficiency Gains

Table 2 presents the average 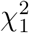 statistics across all loci that reached genome-wide significance (*α* = 5 *×* 10^*−*8^) in the overall sample according to at least one of the association methods. Average 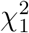 for the asthmatic subgroup are presented in Web Table S6. The empirical *efficiency gain* (O-INT vs. UAT) was defined as the ratio of the non-centrality parameters minus one, where the non-centrality parameters were estimated using loci that reached significance according to at least one association method:

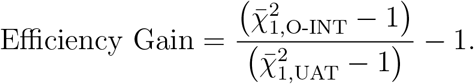

**Table 2.** Empirical Efficiency and Discovery Gains for Lung Function GWAS in the UK Biobank (n = 292K). Genome-wide significance was declared at α = 5 10^*−*8^. The average 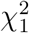 statistics are reported across all loci detected by at least one of the association tests. The empirical efficiency gain, comparing O-INT with UAT, is the ratio of the estimated 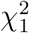 non-centrality parameters minus 1. The counts of significant associations are reported after LD clumping within 1000 kb radii at r^2^ = 0.2 to remove redundant signals. The discovery gain, comparing O-INT with UAT, is the ratio of the number of associations uniquely identified by O-INT to the total number of associations detected.

| Average $\chi^2_1$ | | | | | |
| --- | --- | --- | --- | --- | --- |
| Trait | UAT | D-INT | I-INT | O-INT | Efficiency Gain (%) |
| FEV1 | 56.77 | 57.55 | 64.69 | 63.59 | 12 |
| FVC | 37.14 | 46.91 | 51.47 | 50.59 | 37 |
| FEV1/FVC | 63.21 | 83.76 | 83.00 | 83.63 | 33 |
| lnPEF | 17.72 | 52.44 | 65.40 | 64.11 | 278 |
| Significant Associations |  |  |  |  |  |
| Trait | UAT | D-INT | I-INT | O-INT | Discovery Gain (%) |
| FEV1 | 331 | 352 | 422 | 398 | 15 |
| FVC | 213 | 323 | 375 | 364 | 38 |
| FEV1/FVC | 450 | 653 | 649 | 652 | 28 |
| lnPEF | 39 | 202 | 270 | 251 | 79 |

In all cases the average 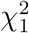 statistics of the INT-based tests exceeded those of UAT, both in the overall analysis and in the asthmatic subgroup. Table 2 also presents the counts of genome-wide significant associations after LD ‘clumping’ to reduce redundant signals. Counts for the asthmatic subgroup are presented in Web Table S6. The empirical *discovery gain* (O-INT vs. UAT) was defined as:

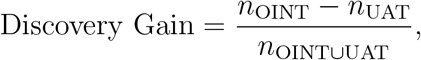

where *n*_OINT_ is the number of associations identified by O-INT only, *n*_UAT_ is the number of associations identified by UAT only (if any), and *n*_OINT*∪*UAT_ is the number of associations identified by either O-INT or UAT. In all cases, the INT-based tests discovered more independent (at *r*^2^ = 0.2) associations with the target phenotype than UAT. All associations that reach genome-wide significance in the asthmatic subgroup, according to either UAT or the INT-based tests, reached significance according to O-INT in the overall sample.

The efficiency and discovery gains were more dramatic for those traits whose residuals were less normally distributed (Web Figure S6). However, the INT-based tests were consistently more powerful than the UAT, even when the normal residual assumption was not unreasonable. Consistent with the simulations, the power of O-INT was always intermediate between that of D-INT and I-INT. For a given trait, the number of discoveries by O-INT was generally closer to the number of discoveries by the more powerful of D-INT and I-INT.

## 5. Discussion

In this paper, we have systematically investigated the utility of different INT-based association tests for GWAS of quantitative traits with non-normally distributed residuals. We formally defined the Direct (D-INT) and Indirect (I-INT) INT-based tests, demonstrating that these approaches are adapted to different underlying data generating processes. D-INT posits that the outcome could have arisen from a monotone transformation of a latent normal trait, whereas I-INT posits that the outcomes has additive but potentially non-normal residuals. When covariate effects are small, the two approaches are approximately equivalent under the null hypothesis of no genetic effect; and in the absence of covariates, the two approaches are identical.

For non-normally distributed quantitative traits, INT-based tests provided nominal control of the type I error by *n* = 10^3^, whereas the UAT exhibited excess type I error even at *n* = 10^5^. Moreover, the INT-based tests were consistently more powerful than UAT. Neither D-INT nor I-INT was uniformly more powerful. To obviate the need for choosing between them, we have proposed an adaptive omnibus test (O-INT). O-INT combines the p-values from D-INT and I-INT via Cauchy combination (Liu and Xie, 2019; Liu et al., 2019), and may easily be extended to incorporate p-values from complementary (e.g. non-parametric) association tests. In simulations and data applications, O-INT provided valid and powerful inference that was robust to the underlying data generating process. As a compromise between complementary methods, O-INT cannot be expected to outperform both D-INT and I-INT. However, the performance of O-INT was similar to the more efficient of the component tests. O-INT was uniformly more powerful than UAT, and is applicable whenever UAT is applicable. All INT-based tests (D-INT, I-INT, O-INT) have been implemented in the R package `RNOmni`, which is available on CRAN. We further demonstrated the utility of INT-based association tests through GWAS of spirometry traits from the UK Biobank.

In D-INT, the INT is applied directly to the phenotype, and the transformed phenotype is regressed on genotype and covariates. Under the complete null hypothesis of no genetic or covariate effects, D-INT is asymptotically exact, and when the covariate effects are small, D-INT holds approximately. I-INT is a two-stage procedure. Different variants of I-INT have been considered in the literature. In all approaches, the phenotype is first regressed on covariates to obtain residuals. In the second stage, the INT-transformed residuals are regressed on genotype, with or without a secondary adjustment for genetic PCs. To provide guidance on which approach to use in practice, we formally derived I-INT, starting from the assumption that the observed phenotype follows a linear regression model with a non-normally distributed residual. Our derivations indicate that, during the second stage of I-INT, the transformed phenotypic residuals should be regressed on genotypic residuals, which are the residuals obtained by regressing genotype on covariates. This second stage is equivalent to regressing the INT-transformed phenotypic residuals on genotype while performing a secondary adjustment for covariates. Therefore, all covariates, including genetic PCs, should be included in both the first and second stage regressions. Under the standard null hypothesis of no genetic effects, I-INT is asymptotically exact, and under the complete null hypothesis of neither genetic nor covariate effects, D-INT and I-INT are asymptotically equivalent.

The use of INT does not compromise the validity of association testing, whose primary objective is to determine whether there is evidence that genotype is associated with the phenotype. Moreover, INT is useful for estimating standardized effect sizes. After INT, any absolutely continuous random variable is unitless, with mean zero, unit variance, and a common limiting distribution. Consequently, effect sizes estimated after INT are comparable across traits measured in different units and along different dimensions. Standardized effect sizes estimated via D-INT (Cade et al., 2016; Abbott et al., 2017) and via I-INT (Kettunen et al., 2012; Consortium et al., 2017) have been reported in numerous applications.

A limitation of INT-based tests is the restriction to absolutely continuous phenotypes. The INT cannot ensure asymptotic normality of a distribution with discrete probability masses. Our simulation studies and data application were restricted to common variants, those having a sample MAF exceeding 5%. For variants with lower MAF, unequal sample sizes can result in non-constant variance across minor allele count strata. This heteroscedasticity is not remedied by INT, and is likely more deleterious than residual non-normality (Beasley et al., 2009). A future direction is to develop set-based tests that leverage the INT to improve power in rare variant association testing.

Finally, this paper has focused on GWAS of independent subjects. However, INT-based tests can be extended to the correlated data setting using linear mixed models (LMMs). We plan to develop INT-based tests for LMMs in which that correlation across related subjects is modeled via a random effect whose covariance pattern depends on the genetic relatedness matrix (Kang et al., 2010; Loh et al., 2015; Chen et al., 2016). A similar modeling strategy can accommodate longitudinal phenotypes arising from repeated measurements on the same subjects across time (Chen et al., 2019).

## Acknowledgments

This work was supported by F31 HL140822 (to Z. M.); by R35 HL135818 (to S. R.); and by R35 CA197449, P01-CA134294, U01-HG009088, U19-CA203654, and R01-HL113338 (to X. L.). We thank the referees for their insightful comments on our manuscript. We would like to thank the Editor, the Associate Editor, and two referees for their helpful comments that have improved this paper.

## Software

The association tests described in this paper (D-INT, I-INT, O-INT) are available in the R package on Github: https://github.com/zrmacc/RNOmni and on CRAN: https://cran.r-project.org/web/packages/RNOmni/index.html. Links to this and related software may also be found at https://content.sph.harvard.edu/xlin/software.html.

