## Supporting Information for "Operating Characteristics of the Rank-Based Inverse Normal Transformation for Quantitative Trait Analysis in Genome-Wide Association Studies"

September 13th, 2019

### Web Table S1

Table 1: **Empirical Type I Error ($\times 10^2$) of INT-based Tests at $\alpha = 0.05$ across $R = 10^8$ Simulation Replicates.** Size simulations were conducted under the $H_0 : \beta_G = 0$ at sample sizes ranging from $n = 10^3$ to $n = 10^5$. The following association tests were evaluated: the untransformed association test (UAT), direct inverse normal transformation (D-INT), indirect INT (I-INT), and omnibus INT (O-INT). Each test was applied to a normal phenotype, a skewed phenotype with $\chi^2_1$ residuals, a kurtotic phenotype with $t_3$ residuals, and a phenotype whose logarithm was normal.

| | | Sample Size | | |
|---|---|---|---|---|
| Phenotype | Test | $n = 10^3$ | $n = 10^4$ | $n = 10^5$ |
| Normal | BAT | 4.99 | 5.00 | 5.00 |
| Normal | D-INT | 4.96 | 4.99 | 5.00 |
| Normal | I-INT | 5.01 | 5.00 | 5.00 |
| Normal | O-INT | 4.99 | 5.00 | 5.00 |
| Skewed | UAT | 4.96 | 5.00 | 5.00 |
| Skewed | D-INT | 4.96 | 5.00 | 5.03 |
| Skewed | I-INT | 4.70 | 4.89 | 4.97 |
| Skewed | O-INT | 4.96 | 5.10 | 5.16 |
| Kurtotic | UAT | 5.02 | 5.02 | 5.01 |
| Kurtotic | D-INT | 4.97 | 5.00 | 5.00 |
| Kurtotic | I-INT | 5.00 | 5.00 | 5.00 |
| Kurtotic | O-INT | 5.00 | 5.02 | 5.02 |
| Log-Normal | UAT | 4.86 | 4.98 | 5.04 |
| Log-Normal | D-INT | 4.97 | 4.99 | 5.00 |
| Log-Normal | I-INT | 4.09 | 4.12 | 4.15 |
| Log-Normal | O-INT | 4.67 | 4.70 | 4.72 |

### Web Table S2

Table 2: **Empirical Type I Error ($\times 10^3$) of INT-based Tests at $\alpha = 10^{-3}$ across $R = 10^8$ Simulation Replicates.** Size simulations were conducted under the $H_0 : \beta_G = 0$ at sample sizes ranging from $n = 10^3$ to $n = 10^5$. The following association tests were evaluated: the untransformed association test (UAT), direct inverse normal transformation (D-INT), indirect INT (I-INT), and omnibus INT (O-INT). Each test was applied to a normal phenotype, a skewed phenotype with $\chi^2_1$ residuals, a kurtotic phenotype with $t_3$ residuals, and a phenotype whose logarithm was normal.

| | | Sample Size | | |
|---|---|---|---|---|
| Phenotype | Test | $n = 10^3$ | $n = 10^4$ | $n = 10^5$ |
| Normal | UAT | 1.00 | 1.00 | 1.00 |
| Normal | D-INT | 0.94 | 1.00 | 1.00 |
| Normal | I-INT | 0.99 | 1.00 | 0.99 |
| Normal | O-INT | 0.97 | 1.00 | 0.99 |
| Skewed | UAT | 1.15 | 1.03 | 1.00 |
| Skewed | D-INT | 0.97 | 1.00 | 1.02 |
| Skewed | I-INT | 0.84 | 0.95 | 0.98 |
| Skewed | O-INT | 0.93 | 1.00 | 1.03 |
| Kurtotic | UAT | 1.21 | 1.11 | 1.05 |
| Kurtotic | D-INT | 0.95 | 1.00 | 1.00 |
| Kurtotic | I-INT | 0.98 | 0.99 | 1.00 |
| Kurtotic | O-INT | 0.97 | 1.00 | 1.00 |
| Log-Normal | UAT | 1.78 | 1.29 | 1.13 |
| Log-Normal | D-INT | 0.94 | 0.99 | 1.00 |
| Log-Normal | I-INT | 0.61 | 0.61 | 0.65 |
| Log-Normal | O-INT | 0.78 | 0.82 | 0.85 |

### Web Table S3

Table 3: **Comparison of Empirical Type I Error ($\times 10^6$) for INT and Nonparametric Testing at $\alpha = 10^{-6}$ across $R = 10^8$ Simulation Replicates.** Size simulations were conducted under the $H_0 : \beta_G = 0$ at sample sizes $n = 10^3$ and $n = 10^4$. To permit direct application of the nonparametric Kruskal-Wallis (KW), there simulations were performed in the absence of covariates. Lacking covariates, the INT-based tests (direct, indirect, and omnibus) are all identical. UAT refers to the untransformed association test. Each test was applied to a normal phenotype, a skewed phenotype with $\chi_1^2$ residuals, a kurtotic phenotype with $t_3$ residuals, and a phenotype whose logarithm was normal.

| | | Sample Size | |
|---|---|---|---|
| Phenotype | Test | $n = 10^3$ | $n = 10^4$ |
| Normal | UAT | 0.77 | 0.99 |
| Normal | INT | 0.78 | 0.98 |
| Normal | KW | 0.38 | 0.76 |
| Skewed | UAT | 4.88 | 1.35 |
| Skewed | INT | 0.64 | 1.07 |
| Skewed | KW | 0.47 | 0.86 |
| Kurtotic | UAT | 7.67 | 3.45 |
| Kurtotic | INT | 0.62 | 0.83 |
| Kurtotic | KW | 0.37 | 0.92 |
| Log-Noraml | UAT | 19.12 | 3.33 |
| Log-Noraml | INT | 0.76 | 1.11 |
| Log-Noraml | KW | 0.37 | 1.03 |

### Web Table S4

Table 4: **LD Score Regression Analysis of Z-scores from Association Testing with Lung Function Traits in the UK Biobank ($n$ = 292K).** LD scores were calculated within the UK Biobank, and LDSC was performed using summary statistics from association testing at all 361K loci with common minor alleles. An LDSC intercept significantly exceeding 1 is indicative of residual confounding. An estimate of the additive heritability $h^2$ explained by loci in the analysis is also provided.

| Trait | Test | Intercept | SE(Int) | $h^2$ | SE($h^2$) |
|---|---|---|---|---|---|
| FEV1 | UAT | 0.84 | 0.02 | 0.24 | 0.01 |
| FEV1 | D-INT | 0.82 | 0.02 | 0.25 | 0.01 |
| FEV1 | I-INT | 0.81 | 0.02 | 0.27 | 0.01 |
| FEV1 | O-INT | 0.81 | 0.02 | 0.27 | 0.01 |
| FVC | UAT | 0.86 | 0.02 | 0.21 | 0.01 |
| FVC | D-INT | 0.83 | 0.02 | 0.25 | 0.01 |
| FVC | I-INT | 0.81 | 0.02 | 0.28 | 0.01 |
| FVC | O-INT | 0.81 | 0.02 | 0.27 | 0.01 |
| FEV1/FVC | UAT | 0.86 | 0.02 | 0.22 | 0.01 |
| FEV1/FVC | D-INT | 0.86 | 0.02 | 0.27 | 0.01 |
| FEV1/FVC | I-INT | 0.84 | 0.02 | 0.28 | 0.01 |
| FEV1/FVC | O-INT | 0.84 | 0.02 | 0.28 | 0.01 |
| lnPEF | UAT | 0.95 | 0.01 | 0.06 | 0.004 |
| lnPEF | D-INT | 0.88 | 0.02 | 0.16 | 0.01 |
| lnPEF | I-INT | 0.86 | 0.02 | 0.18 | 0.01 |
| lnPEF | O-INT | 0.86 | 0.02 | 0.18 | 0.01 |

### Web Table S5

Table 5: **LD Score Regression Analysis of Z-scores from Association Testing with Lung Function Traits in the Asthmatic Subgroup ($n$ = 29K).** LD scores were calculated within the UK Biobank, and LDSC was performed using summary statistics from association testing at all 361K loci with common minor alleles. An LDSC intercept significantly exceeding 1 is indicative of residual confounding. An estimate of the additive heritability $h^2$ explained by loci in the analysis is also provided.

| Trait | Test | Intercept | SE(Int) | $h^2$ | SE($h^2$) |
|---|---|---|---|---|---|
| FEV1 | UAT | 0.97 | 0.01 | 0.29 | 0.03 |
| FEV1 | D-INT | 0.96 | 0.01 | 0.33 | 0.04 |
| FEV1 | I-INT | 0.95 | 0.01 | 0.36 | 0.04 |
| FEV1 | O-INT | 0.95 | 0.01 | 0.35 | 0.04 |
| FVC | UAT | 0.98 | 0.01 | 0.18 | 0.03 |
| FVC | D-INT | 0.96 | 0.01 | 0.30 | 0.03 |
| FVC | I-INT | 0.96 | 0.01 | 0.34 | 0.03 |
| FVC | O-INT | 0.95 | 0.01 | 0.33 | 0.03 |
| FEV1/FVC | UAT | 0.99 | 0.01 | 0.29 | 0.04 |
| FEV1/FVC | D-INT | 0.98 | 0.01 | 0.33 | 0.04 |
| FEV1/FVC | I-INT | 0.98 | 0.01 | 0.31 | 0.04 |
| FEV1/FVC | O-INT | 0.98 | 0.01 | 0.32 | 0.04 |
| lnPEF | UAT | 1.01 | 0.01 | 0.06 | 0.02 |
| lnPEF | D-INT | 0.97 | 0.01 | 0.24 | 0.03 |
| lnPEF | I-INT | 0.96 | 0.01 | 0.27 | 0.04 |
| lnPEF | O-INT | 0.96 | 0.01 | 0.26 | 0.03 |

### Web Table S6

Table 6: **Empirical Efficiency and Discovery Gains for Lung Function GWAS in the Asthmatic Subgroup ($n$ = 29K).** Genome-wide significance was declared at $\alpha = 5 \times 10^{-8}$. The average $\chi_1^2$ statistics are reported across all loci detected by at least one of the association tests. The empirical efficiency gain, comparing O-INT with UAT, the ratio of the estimated $\chi_1^2$ non-centrality parameters minus 1. The counts of significant associations are reported after LD clumping within 1000 kb radii at $r^2 = 0.2$ to remove redundant signals. The discovery gain, comparing O-INT with UAT, is the ratio of the number of associations uniquely identified by O-INT to the total number of associations detected.

| **Average $\chi_1^2$** | | | | | |
|---|---|---|---|---|---|
| Trait | UAT | D-INT | I-INT | O-INT | Efficiency Gain (%) |
| FEV1 | 27.56 | 31.36 | 29.84 | 31.06 | 13 |
| FVC | 20.96 | 38.50 | 37.84 | 38.35 | 87 |
| FEV1/FVC | 50.76 | 53.60 | 53.10 | 53.52 | 6 |
| lnPEF | 14.22 | 38.54 | 44.75 | 43.57 | 222 |
| **Significant Associations** | | | | | |
| Trait | UAT | D-INT | I-INT | O-INT | Discovery Gain (%) |
| FEV1 | 3 | 6 | 4 | 5 | 40 |
| FVC | 0 | 4 | 3 | 4 | 100 |
| FEV1/FVC | 14 | 15 | 15 | 16 | 12 |
| lnPEF | 0 | 3 | 3 | 3 | 100 |

### Web Figure S1

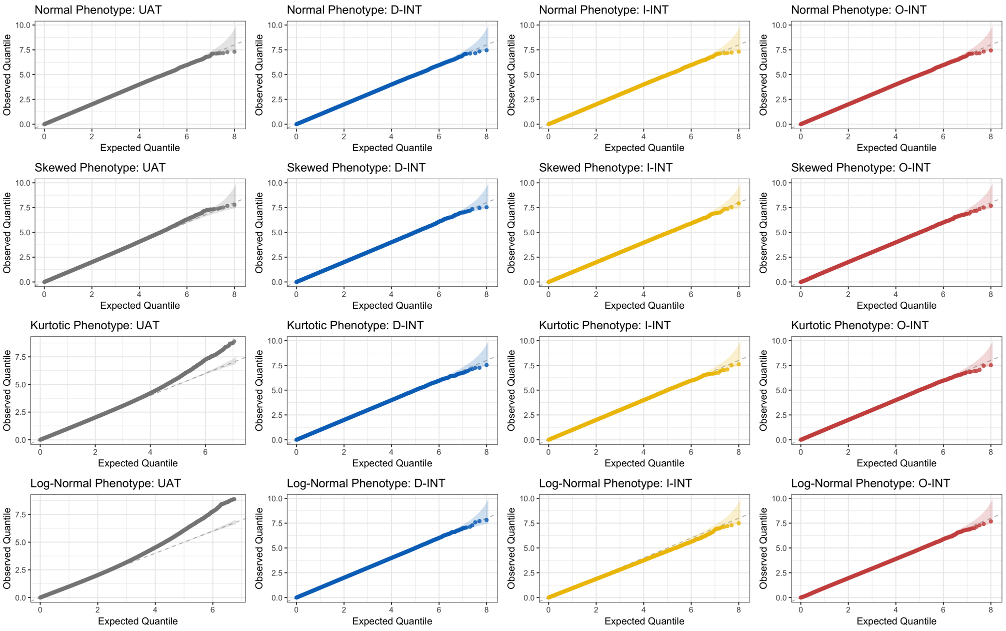


Figure 1: **Distribution of Association p-values Under the Null at $n = 10^4$ across $R = 10^8$ Simulation Replicates.** Rows correspond to different phenotype distributions. The first is normal, the second skewed ($\chi_1^2$), the third kurtotic ($t_3$), and the fourth log-normal. Columns correspond to different association tests. The first is the untransformed association test (UAT), the second is the direct inverse normal transformation (D-INT), the third is indirect INT (I-INT), and the fourth column is omnibus INT (O-INT).

### Web Figure S2

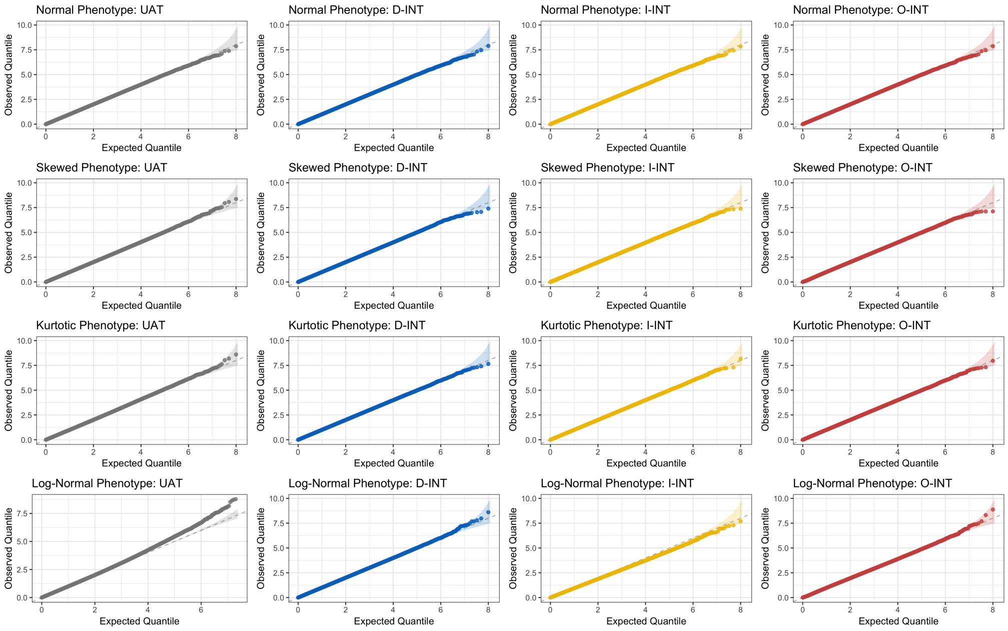


Figure 2: **Distribution of Association p-values Under the Null at $n = 10^5$ across $R = 10^8$ Simulation Replicates.** Rows correspond to different phenotype distributions. The first is normal, the second skewed ($\chi_1^2$), the third kurtotic ($t_3$), and the fourth log-normal. Columns correspond to different association tests. The first is the untransformed association test (UAT), the second is the direct inverse normal transformation (D-INT), the third is indirect INT (I-INT), and the fourth column is omnibus INT (O-INT).

### Web Figure S3

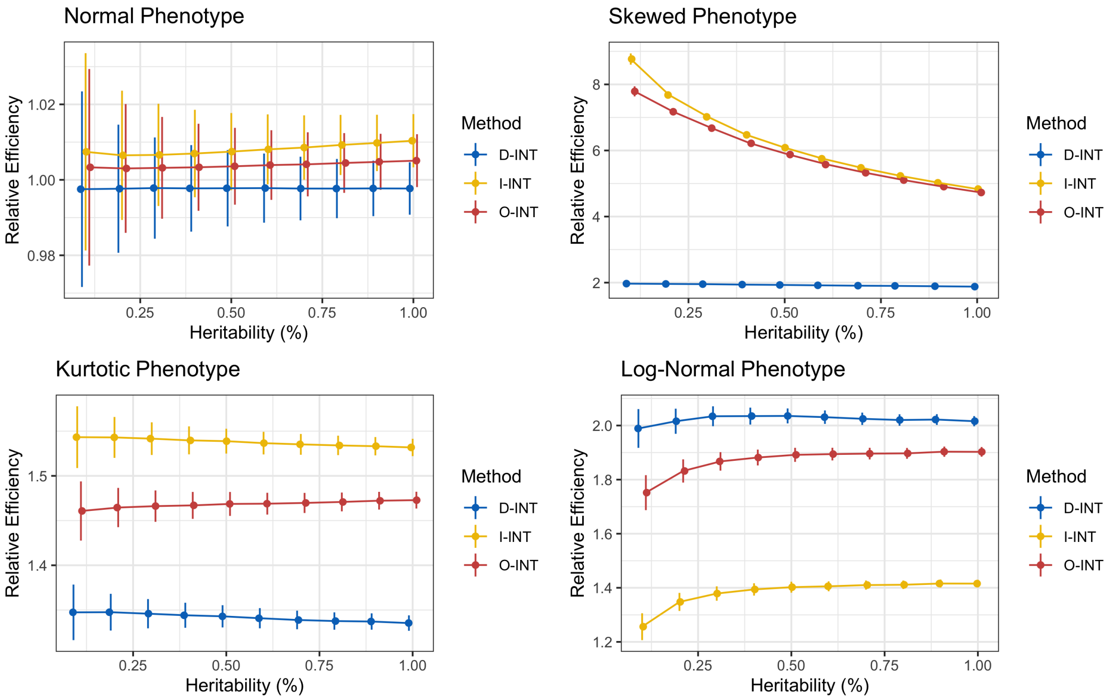


Figure 3: **Relative Efficiency (RE) Curves across $\boldsymbol{R} = \mathbf{10^6}$ Simulation Replicates.** RE was calculated as the ratio of $\chi^2_1$ non-centrality parameters. This metric has the advantage of depending on neither $\alpha$ level nor sample size $n$. Simulations were conducted at heritabilities ranging from 0.1% and 1.0%. Gray is the UAT, blue the D-INT, yellow the I-INT, red the O-INT. The first phenotype has normal residuals; the second has $\chi^2_1$ residuals; the third phenotype has $t_3$ residuals; and the log of the fourth phenotype has normal residuals.

### Web Figure S4

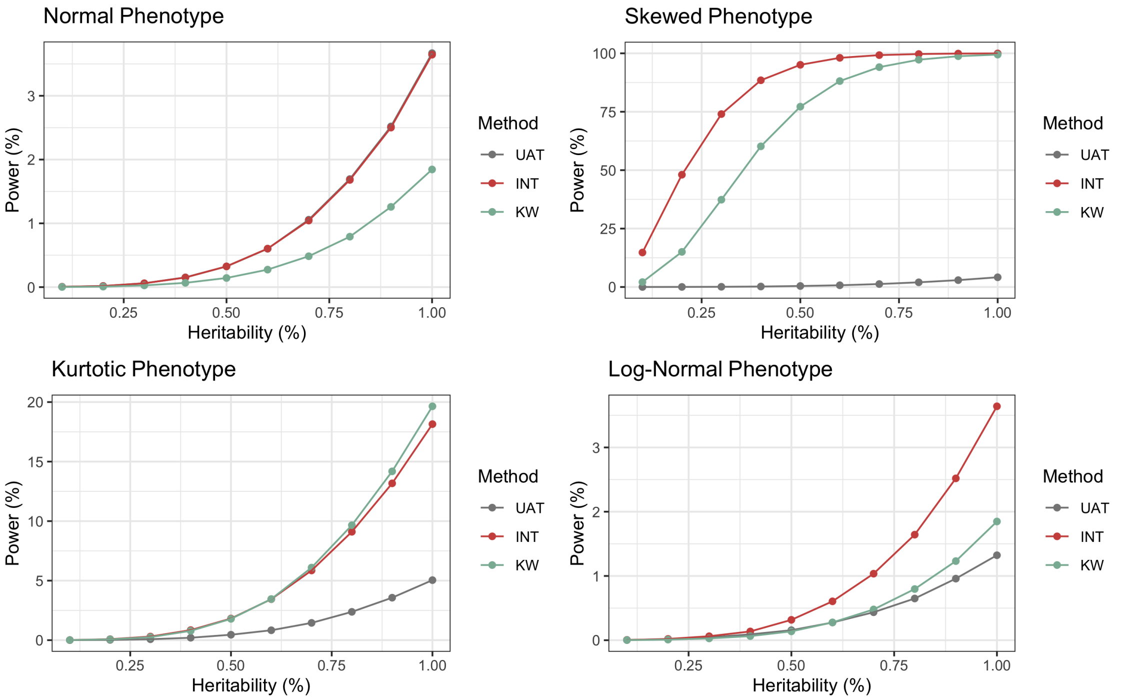


Figure 4: **Power Curves Comparing INT-based Testing with the Nonparametric Kruskal-Wallis (KW) Test at $n = 10^3$ across $R = 5 \times 10^5$ Simulation Replicates**. To permit direct application of the KW test, these simulations were conducted in the absence of covariates. Lacking covariates, the INT-based tests (direct, indirect, and omnibus) are all identical. The heritability, or proportion of phenotype variation explained by genotype, was varied between 0.1% and 1.0%. The untransformed association test (UAT) is displayed for reference.

### Web Figure S5

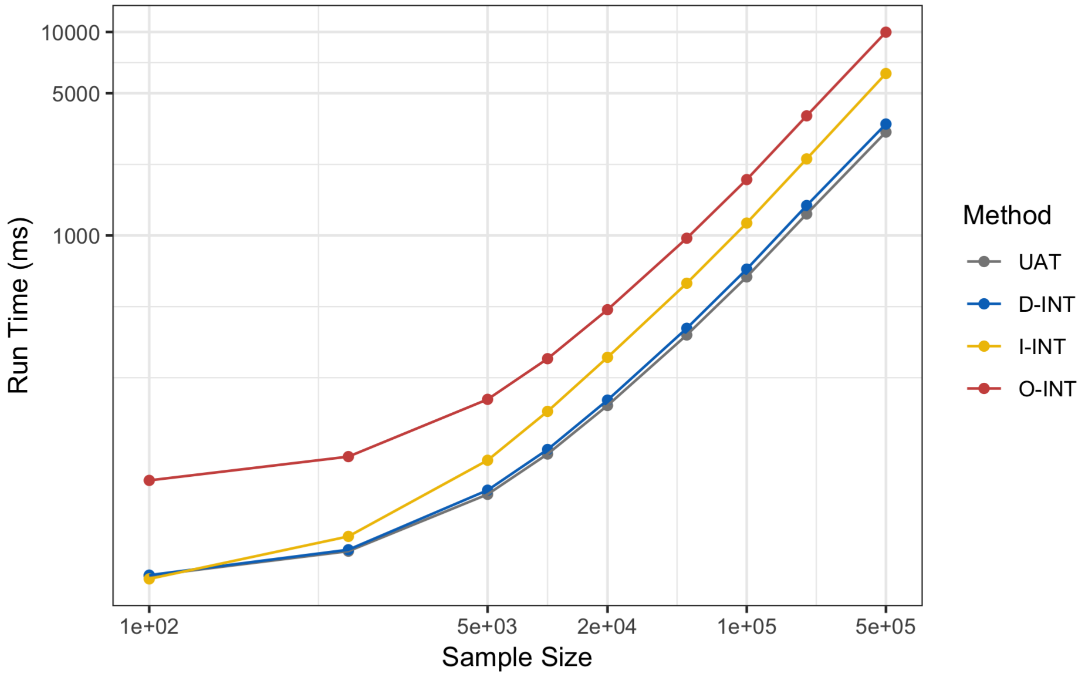


Figure 5: **Median Run Time for $10^2$ Association Tests across $R = 10^3$ Simulation Replicates as a Function of Sample Size $n$.** The computational cost of UAT, which is based on standard linear regression, scales linearly in sample size. These results demonstrate that the computational costs of the INT-based association tests scale at the same rate.

#### Web Figure S6

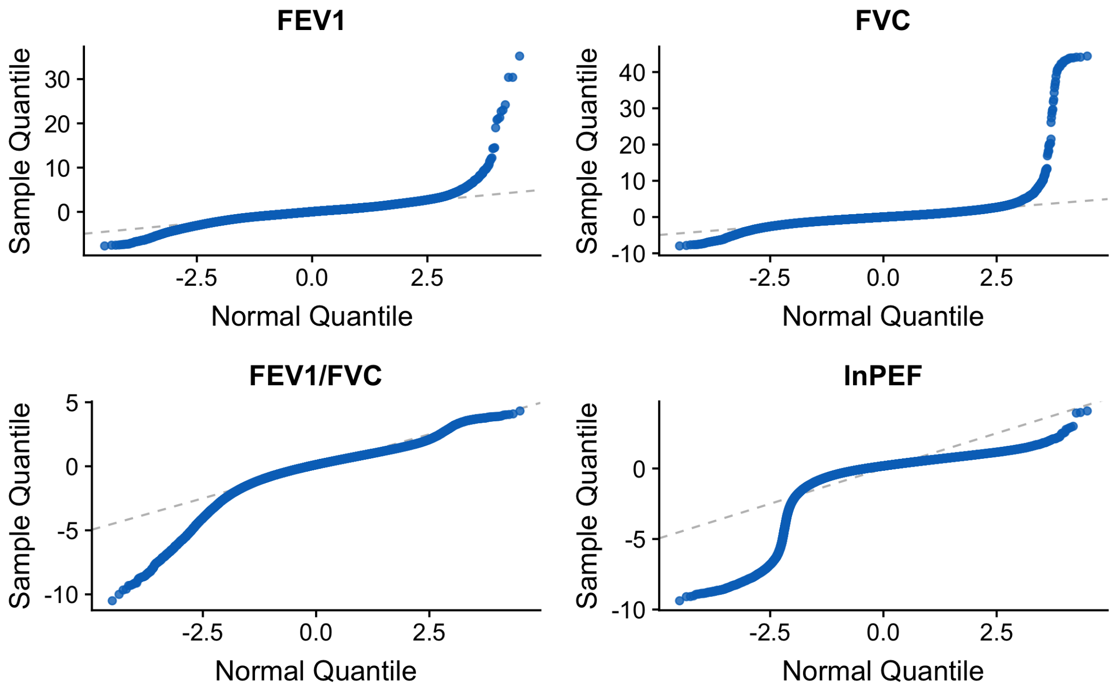


Figure 6: **Normal Quantile-Quantile Plots for Spirometry Traits from the UK Biobank ($n = 292$K).** Each trait was regressed on covariates, including age, sex, height, genotyping array, and genetic PCs, to obtain phenotypic residuals. Residual Z-scores (vertical) are compared with the quantiles of the standard normal distribution (horizontal). Departure from the diagonal indicates non-normality of the residual distribution.

### Web Figure S7

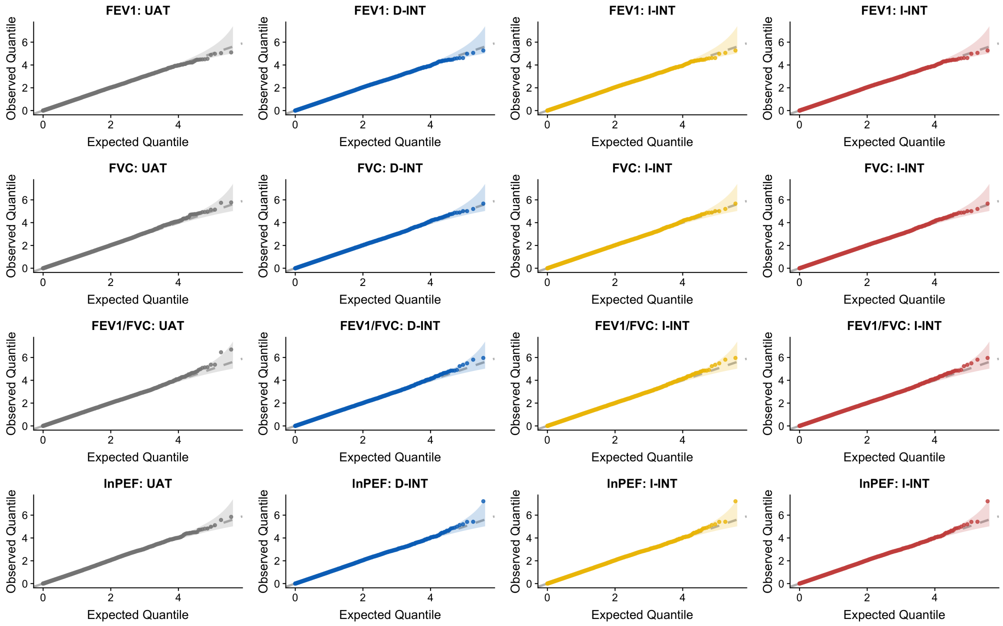


Figure 7: **Distribution of p-values from Association Testing in the UK Biobank after Permuting Genotypic Residuals.** The sample size was $n = 292$K. Genotypes at 361K loci with common minor alleles were included. Genotypic and phenotypic residuals were obtained by first regressing out the effects of covariates, including age, sex, height, genotyping array, and genetic PCs. First column is the untransformed association test (UAT); second column is the direct inverse normal transformation (D-INT); third column is indirect INT (I-INT); fourth column is omnibus INT (O-INT). First row is forced expiratory volume (FEV1); second row is forced vital capacity (FVC); third row is the FEV1/FVC ratio; fourth row is the logarithm of peak expiratory flow (lnPEF).
